# Chronic Unpredictable Stress Drives Non-pathological, Adaptive Metabolic Hormone Reprogramming with Reduced Insulin Resistance Linked to Depressive-like Behaviors

**DOI:** 10.64898/2026.02.05.703942

**Authors:** Yueyang Luo, Tangcong Chen, Mengdie Li, Marta Kubera, Yingqian Zhang, Yulan Huang, Michael Maes

## Abstract

Major Depressive Disorder (MDD) is increasingly conceptualized as a neuroimmune-metabolic-oxidative stress (NIMETOX) condition, partly driven by environmental stressors. While new antidepressant strategies have emerged to target NIMETOX pathways, the mechanism by which chronic stress reshapes metabolic regulation-particularly in the absence of clinically overt metabolic disease-remain poorly understood.

Using a 6-week chronic unpredictable mild stress (CUMS) paradigm in metabolically healthy male and female mice, we evaluated depressive-like behaviors alongside circulating glucose and metabolic hormones, including insulin, resistin, Glucose-dependent Insulinotropic Polypeptide (GIP), Glucagon-like Peptide-1 (GLP-1), glucagon, ghrelin, leptin, and Plasminogen Activator Inhibitor-1 (PAI-1). We further assessed whether simvastatin, curcumin, S-adenosyl methionine (SAMe), or pyrrolidine dithiocarbamate (PDTC), an NF-κB inhibitor, normalize CUMS-induced metabolic and behavioral alterations, using fluoxetine as a reference antidepressant.

CUMS induced a hypoinsulinemic state accompanied by enhanced basal insulin sensitivity and reduced insulin resistance, while central appetite regulation and adipose inflammatory profiles remained preserved. Indices of lower insulin resistance were strongly associated with sucrose preference and immobility time. Glucagon and PAI-1 showed positive associations with novel object recognition performance, whereas GIP and leptin were inversely related to open-field activity. Glucose levels correlated positively with rearing behavior. Despite significant improvements in depressive-like behaviors, none of the pharmacological interventions normalized the stress-induced metabolic hormone profile. Marked sex differences were observed, with females displaying a catabolic, immune-ready phenotype and males showing a relative anabolic bias.

These findings indicate that CUMS induces a non-pathological, adaptive metabolic hormone reprogramming rather than metabolic dysfunction, supporting the interpretation of stress-related metabolic changes as resilient adaptations within the NIMETOX framework.

## 1. Introduction

Major depressive disorder (MDD) affects over 280 million people worldwide and is often chronic or relapsing (Hasin et al., 2018). It causes substantial disability and premature mortality, including through suicide and cardiometabolic comorbidity (Haagsma et al., 2016). Increasingly, MDD is viewed as a multisystem disorder spanning neuroimmune–metabolic and oxidative-stress pathways (NIMETOX) (Maes et al., 2025), with reproducible alterations in inflammation, energy homeostasis, and stress-system regulation across clinical and preclinical studies (López-López et al., 2016; Maes et al., 2025). This supports biomarker-informed, precision approaches to antidepressant treatment.

Within the NIMETOX framework, metabolic syndrome (MetS) is not only a frequent comorbidity but also a clinically salient correlate and potential contributor to depressive phenotypes (Maes et al., 2025). Epidemiological and longitudinal studies support bidirectional associations between depression and MetS-related traits, including impaired fasting glucose and dyslipidemia, although lifestyle factors and medication remain important confounders (Dregan et al., 2020; Moreira et al., 2017, 2019). Mechanistically, a sustained pro-inflammatory milieu—including M1 macrophage activation and elevated IL-1β, IL-6, and TNF-α—can impair insulin signaling and promote peripheral and central insulin resistance, with downstream consequences for glucose homeostasis and neural plasticity (Olefsky & Glass, 2010).

Beyond the glucose–insulin axis, gut-derived incretins and adipose-derived signals intersect with metabolism, immune tone, and behavior. Glucagon-like peptide (GLP-1) and glucose-dependent insulinotropic polypeptide (GIP) regulate insulin/glucagon secretion, satiety, and energy balance (Fasshauer & Blüher, 2015; Kieffer & Francis Habener, 1999). Although clinical evidence remains mixed, GLP-1 receptor agonists show neuroprotective and antidepressant-like effects in preclinical and early clinical studies (Chen et al., 2024). Leptin integrates adiposity and inflammatory status to modulate reward and motivational circuitry, whereas ghrelin links energy deficit and stress responses to feeding and mood (Gajewska et al., 2023; Hontecilla-Prieto et al., 2025). Inflammation-linked mediators such as plasminogen activator inhibitor-1 (PAI-1) and resistin reflect metaflammation and have been associated with vascular risk and cognitive/affective change (Chouchani & Kajimura, 2019; Hotamisligil, 2017). Together, these hormones and adipokines provide a multi-analyte window into neurobiological substrates relevant to anhedonia, anxiety, and recognition memory.

The chronic unpredictable mild stress (CUMS) paradigm is widely used to investigate links among environmental stress, depressive-like behaviors, and NIMETOX pathways (Kubera et al., 2011; Sharma et al., 2024). CUMS reliably induces anhedonia, increased forced-swim immobility, anxiety-like behavior, and recognition memory deficits, accompanied by shifts in inflammatory tone, metabolic indices, and neuroprotective signaling (Antoniuk et al., 2019). This model therefore enables joint characterization of behavioral phenotypes and peripheral biomarkers under sustained stress exposure. Sex is a critical biological variable in depression and metabolism: women show higher lifetime risk and distinct endocrine/inflammatory profiles, and sex differences in stress reactivity, metabolic hormones, and treatment response are well documented (Kuehner, 2017; Slavich & Sacher, 2019). Accordingly, modeling sex as an effect modifier can improve interpretability and translational relevance in preclinical studies.

Current evidence indicates that selective serotonin reuptake inhibitors (SSRIs) such as fluoxetine are inadequate for a substantial proportion of individuals with MDD, with many failing to achieve remission and showing persistent NIMETOX abnormalities (Maes et al., 2012). Consequently, novel and repurposed agents are increasingly explored to complement monoaminergic therapies by targeting NIMETOX pathways that SSRIs do not address (Maes et al., 2012). Candidate augmentation compounds include S-adenosylmethionine (SAMe), a methyl donor that enhances monoamine synthesis and modulates epigenetic regulation (Lu & Mato, 2012); simvastatin, which exerts immunomodulatory and anti-inflammatory effects beyond lipid lowering (Menze et al., 2021); curcumin, a polyphenol with antioxidant and anti-inflammatory properties and antidepressant-like effects in preclinical models (Fan et al., 2018; Yuan et al., 2025); and nuclear factor kappa-light-chain-enhancer of activated B cells (NF-κB) pathway inhibitors such as pyrrolidine dithiocarbamate (PDTC), a thiol-containing antioxidant that suppresses NF-κB signaling (Yin et al., 2015).

However, few studies using the CUMS model have simultaneously profiled indices of insulin sensitivity and glucose homeostasis in conjunction with metabolic hormones, inflammatory adipokines, and depressive-like behaviors. Moreover, explicit testing of sex differences and direct comparison of mechanistically distinct interventions—such as those targeting monoaminergic, epigenetic, lipid, and NF-κB pathways—within a single experimental framework are rare.

Hence, the aims of this study are to (i) characterize how CUMS reprograms the metabolic–affective axis under metabolically uncompromised conditions and quantify baseline sex differences in circulating metabolic hormones; (ii) compare the efficacy of fluoxetine, SAMe, simvastatin, curcumin, and PDTC in modulating serum metabolic profiles in association with depression-like behaviors.

## 2. Materials and methods

### 2.1 Animals and Chronic Unpredictable Mild Stress Model

Eighty-four 3-week-old C57BL/6N mice (sex-balanced) were randomly assigned to control or chronic unpredictable mild stress (CUMS) conditions. Mice were housed under standard conditions (12: 12 h light/dark cycle, 23 ± 1°C) with ad libitum chow and water. The CUMS procedure lasted 6 weeks and consisted of two mild stressors per day applied in a randomized schedule to minimize habituation (see **ESF Table S1**). The experimental timeline is shown in Extended **ESF Fig. 1A**. All procedures were approved by the Institutional Animal Care and Use Committee of Sichuan Junhui Biotechnology Co., Ltd. (IACUC-202411-4-001).

**Figure 1.**
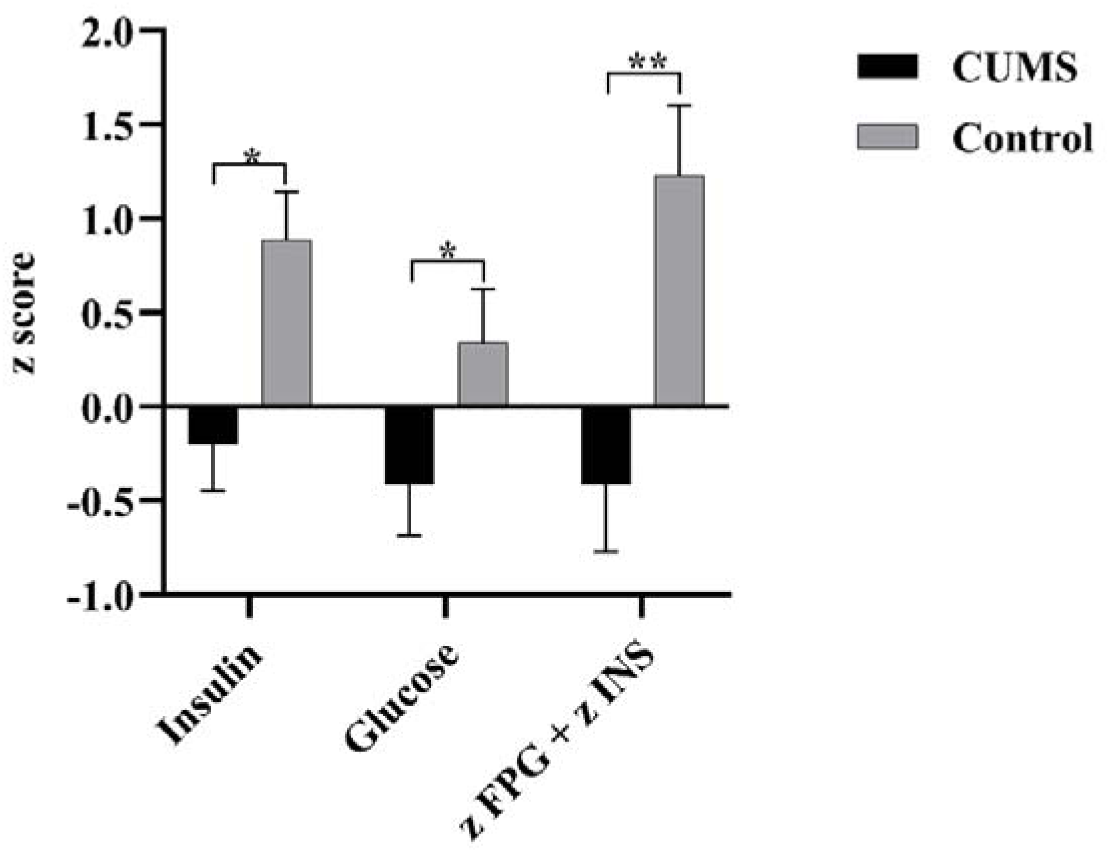
Results of general linear models (GLM) showing sex-adjusted differences in insulin resistance indicators between CUMS and Control. Bars show adjusted estimated marginal means (EMMs) ± standard errors (SE), Error bars represent SE. *, *P* < 0.05; **, *P* < 0.01; ***, *P* < 0.001. CUMS: chronic unpredictable mild stress, FPG: fasting plasma glucose, INS: insulin.

### 2.2 Pharmacological Interventions

Beginning in experimental week 3, mice were randomized into seven groups (n = 12/group, sex-balanced): Control + Vehicle; CUMS + Vehicle; CUMS + Fluoxetine; CUMS + Simvastatin; CUMS + Curcumin; CUMS + SAMe; and CUMS + PDTC. Treatments were administered by gavage once daily for 28 days (from weeks 3 to 6). Drug preparation, dosing, vehicles, and administration routes are provided in **ESF Table S2.**

### 2.3 Body Weight Monitoring and Behavioral Tests

Body weight and food intake were recorded weekly. Behavioral testing was conducted in week 7 in the following order: open-field test (OFT), sucrose preference test (SPT), novel object recognition (NOR), and forced swimming test (FST), with experimenters blinded to group allocation. Primary behavioral outcomes were sucrose preference (SPT), immobility time (FST), locomotor and center activity (OFT), and recognition index (NOR). Full procedures and scoring criteria are described in **ESF Methods S1**.

### 2.4 Serum collection and assays

At least 24 h after the final behavioral test, mice were fasted overnight (∼16 h) and euthanized. Blood was collected by cardiac puncture, and serum was isolated, aliquoted, and stored at −80°C until batch analysis. Eight metabolic hormones (ghrelin, GIP, GLP-1, glucagon, insulin, leptin, PAI-1, resistin) were quantified using a multiplex bead-based assay (Luminex platform) in technical duplicates. Fasting serum glucose was measured using a colorimetric assay. Detailed protocols and instrument settings are provided in **ESF Methods S2**.

### 2.5 Statistics

Analyses were performed in SPSS 27.0 (two-tailed, α = 0.05). Sex was included as a fixed factor where applicable. Group differences in biomarker panels were evaluated using multivariate analysis of covariance (MANCOVA), and behavioral outcomes were analyzed using unpaired t-tests (two-group) or one-way ANOVA (multi-group) with appropriate post hoc comparisons. A composite fasting glycemic/insulinemic burden index was computed as zFPG+zINS. Exploratory regression analyses relating biomarkers to behavioral outcomes are described in **ESF Methods S3**. One control female died before the endpoint and was excluded; all other data were complete.

## 3. Results

### 3.1 Validation of the CUMS Model

Behavioral tests confirmed the induction of depression-like phenotypes following CUMS exposure. As shown in **ESF Fig. 1B**, CUMS mice showed a significant decrease in sucrose preference compared to controls. This deficit was significantly improved by treatment with fluoxetine, simvastatin, and curcumin. Similarly, in the FST, CUMS significantly increased immobility time (**ESF Fig. 1C**). This increase in immobility time was significantly attenuated by fluoxetine, simvastatin, and curcumin. The efficacy of fluoxetine validated the model’s pharmacological responsiveness.

In order to examine the effects of CUMS, drugs and sex on the metabolic hormone data and glucose levels, we employed a multivariate general linear model (GLM; MANOVA) with fixed effects of treatment (control, CUMS, and the different drugs), as well as sex. As such, we compared n = 42 males and n = 41 females.

### 3.2 Effects of CUMS on Metabolic Profiles

**Figure 1** shows sex-adjusted group comparisons between CUMS and Control for fasting insulin, fasting glucose, and the composite insulin resistance index (z FPG + z INS). Compared with Controls, the CUMS group exhibited significantly lower insulin and glucose. Furthermore, the z FPG + z INS was significantly lower in the CUMS group.

### 3.3 Sex Differences in Metabolic Hormone Levels

**#Figure 2** displays treatment-adjusted sex differences across metabolic hormones. Females showed significantly higher resistin, PAI-1, glucagon, and ghrelin than males. In contrast, males exhibited significantly higher leptin and insulin. No significant sex differences were observed for GIP, GLP-1, or glucose.

**Figure 2.**
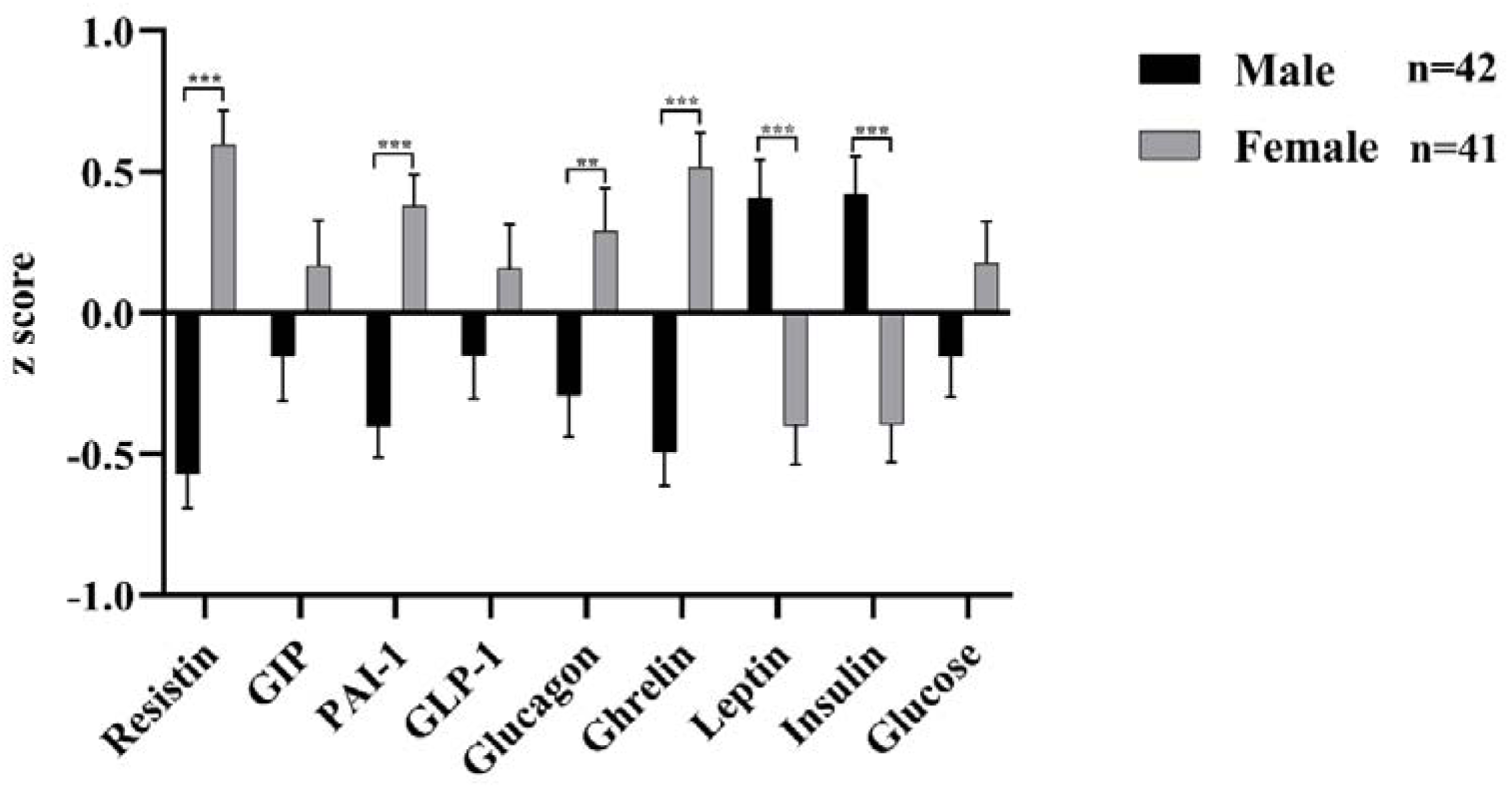
Results of general linear models (GLM) showing treatment-adjusted differences in metabolic indicators between males and females. Bars show adjusted estimated marginal means (EMMs) ± standard errors (SE), Error bars represent SE. *, *P* < 0.05; **, *P* < 0.01; ***, *P* < 0.001. GIP: gastric inhibitory polypeptide, PAI-1: plasminogen activator inhibitor-1, GLP-1: glucagon-like peptide-1

### 3.4 Effects of Pharmacological Treatments on Metabolic Parameters

**Figure 3** presents the metabolic profile across different treatment groups. Adipokine analysis revealed that SAMe significantly elevated resistin levels relative to the CUMS group, while PDTC suppressed them compared to the control. Leptin levels remained unchanged. For gastrointestinal hormones, ghrelin levels were significantly altered: curcumin caused a mild elevation, SAMe a moderate elevation, and PDTC a mild-to-moderate elevation compared to CUMS and control groups, respectively. GIP and glucagon showed no significant variations. Assessments of pancreatic function and glucose homeostasis indicated profound suppression of insulin secretion in the CUMS and all treatment groups compared with controls. Simvastatin significantly increased GLP-1 levels compared with the CUMS group. Curcumin treatment led to a pronounced increase in PAI-1, exceeding levels in the CUMS group and most other groups. Finally, both SAMe and PDTC treatments resulted in significantly higher glucose levels than in the CUMS group.

**Figure 3.**
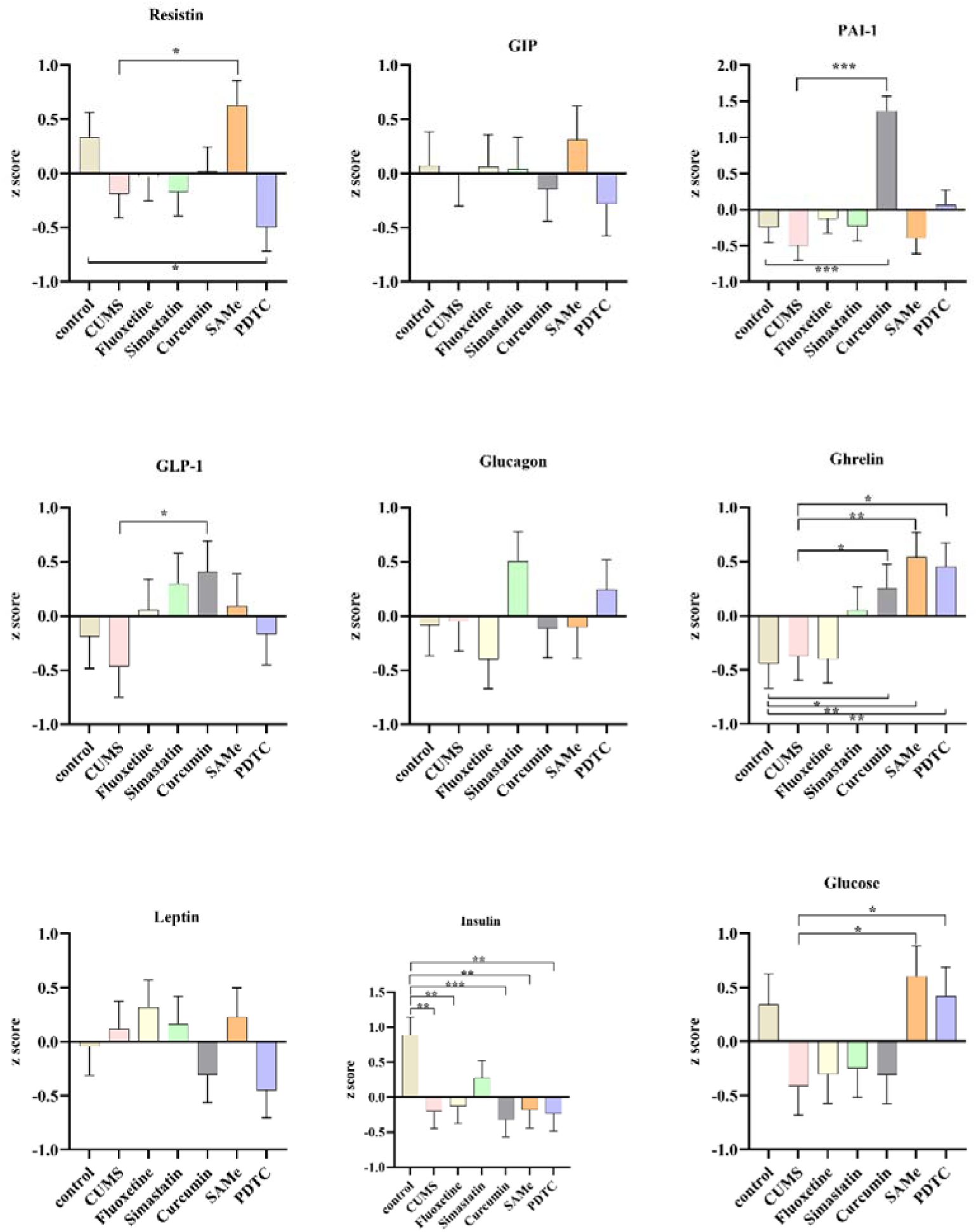
Treatment-related profiles of metabolic indicators: adjusted estimated marginal means (±SE) from MANCOVA Bars show adjusted estimated marginal means (EMMs) ± standard errors (SE) for nine metabolic indicators (Resistin, GIP, PAI-1, GLP-1, Glucagon, Ghrelin, Leptin, Insulin, and Glucose) across treatment groups (Control, CUMS, Fluoxetine, Simvastatin, Curcumin, SAMe, and PDTC). Values are derived from a multivariate analysis of covariance (MANCOVA) with treatment as the fixed factor. Error bars represent SE. *, P < 0.05; **, P < 0.01; ***, P < 0.001. GIP: gastric inhibitory polypeptide, PAI-1: plasminogen activator inhibitor-1, GLP-1: glucagon-like peptide-1, CUMS: chronic unpredictable mild stress, SAMe: S-Adenosylmethionine, PDTC: pyrrolidinedithiocarbamic acid.

### 3.5 Predictors of Metabolic Dysregulation

#### 3.5.1 Models Including Treatments and CUMS Exposure

**Table 1** summarizes the results of linear regression analyses examining the effects of treatment conditions, sex, CUMS exposure, and metabolic hormone levels on behavioral measures. Regression model #1 shows that 21.1% of the variance in the SPT was explained by the z FPG + z INS score and fluoxetine treatment (positive associations), together with CUMS exposure (negative association). Regression model #2 demonstrates that 81.8% of the variance in BW was explained by sex, the z FPG + z INS score, curcumin treatment, and leptin levels (all positive associations). Regression model #3 indicates that 38.2% of the variance in FI was explained by sex and fluoxetine treatment (both positive associations). Regression model #4 shows that 15.0% of the variance in OFT total distance was explained by PDTC treatment (positive association). Regression model #5 demonstrates that 26.6% of the variance in OFT surround distance was explained by PDTC, fluoxetine, and simvastatin treatments (all positive associations). Regression model #6 indicates that 11.1% of the variance in the NOR Preference Index was explained by glucagon and PAI-1 levels (both positive associations). Regression model #7 shows that 12.6% of the variance in FST Immobility Time was explained by the z FPG + z INS score and fluoxetine treatment (both negative associations).

**Table 1.** Linear regression with behavioral measures as dependent variables and treatment, sex, CUMS exposure, and metabolic hormone levels as explanatory variables.

| Dependent Variables | Explanatory Variables | Coefficient statistics |  |  | Model statistics |  |  |  |
| --- | --- | --- | --- | --- | --- | --- | --- | --- |
| | | $\beta$ | t | p | R<sup>2</sup> | F | df | p |
| #1. SPT | Model |  |  |  | 0.211 | 6.951 | 3/78 | < 0.001 |
|  | z FPG + z INS | 0.272 | 2.512 | 0.014 |  |  |  |  |
|  | Fluoxetine | 0.328 | 3.212 | 0.002 |  |  |  |  |
|  | CUMS | -0.190 | -1.752 | 0.084 |  |  |  |  |
| #2. BW | Model |  |  |  | 0.818 | 86.409 | 4/77 | < 0.001 |
|  | Sex | 0.774 | 14.455 | < 0.001 |  |  |  |  |
|  | z FPG + z INS | 0.199 | 3.759 | < 0.001 |  |  |  |  |
|  | Curcumin | 0.147 | 2.946 | 0.004 |  |  |  |  |
|  | Leptin | 0.128 | 2.282 | 0.025 |  |  |  |  |
| #3. FI | Model |  |  |  | 0.382 | 24.371 | 2/79 | < 0.001 |
|  | Sex | 0.586 | 6.619 | < 0.001 |  |  |  |  |
|  | Fluoxetine | 0.196 | 2.220 | 0.029 |  |  |  |  |
| #4. OFT total | Model |  |  |  | 0.150 | 14.171 | 1/80 | < 0.001 |
|  | PDTC | 0.388 | 3.764 | < 0.001 |  |  |  |  |
| #5. OFT surround | Model |  |  |  | 0.266 | 9.429 | 3/78 | < 0.001 |
|  | PDTC | 0.493 | 4.901 | < 0.001 |  |  |  |  |
|  | Fluoxetine | 0.267 | 2.649 | 0.010 |  |  |  |  |
|  | Simvastatin | 0.248 | 2.464 | 0.016 |  |  |  |  |
| #6. NOR Preference Index | Model |  |  |  | 0.111 | 4.824 | 2/77 | 0.022 |
|  | Glucagon | 0.233 | 2.159 | 0.034 |  |  |  |  |
|  | PAI-1 | 0.216 | 1.996 | 0.049 |  |  |  |  |
| #7. FST Immobility Time | Model |  |  |  | 0.126 | 5.681 | 2/79 | 0.005 |
|  | z FPG + z INS | -0.273 | -2.574 | 0.012 |  |  |  |  |
|  | Fluoxetine | -0.265 | -2.500 | 0.014 |  |  |  |  |

SPT: Sucrose preference test. BW: Body weight. FI: Food intake. OFT total: total distance travelled in the Open Field Test.
OFT surround: distance travelled in the surrounding zone of the open field. NOR Preference Index : Novel Object
Recognition Preference Index for the novel object [novel/(novel+familiar)×100%]. FST Immobility Time: Forced Swim Test
Immobility time.

#### 3.5.2 Models with Sex and Metabolic Hormones Only

**Table 2** summarizes the results of linear regression analyses examining the effects of sex and metabolic hormone levels on behavioral measures. Regression model #1 shows that 14.0% of the variance in SPT was explained by the z FPG + z INS score (positive association) and glucagon levels (negative association). Regression model #2 demonstrates that 79.9% of the variance in BW was explained by sex, the z FPG + z INS score, and PAI-1 levels (all positive associations). Regression model #3 indicates that 37.5% of the variance in FI was explained by sex and leptin levels (both positive associations). Regression model #4 shows that 9.9% of the variance in OFT surround distance was explained by GIP and leptin levels (both negative associations). Regression model #5 demonstrates that 10.4% of the variance in OFT rearing was explained by glucose levels (positive association). Regression model #6 indicates that 11.1% of the variance in NOR Preference Index was explained by glucagon and PAI-1 levels (both positive associations). Regression model #7 shows that 5.7% of the variance in FST Immobility Time was explained by the z FPG + z INS score (negative association).

**Table 2.** Linear regression with behavioral measures as dependent variables, sex and metabolic hormone levels as explanatory variables.

| Dependent Variables | Explanatory Variables | Coefficient statistics |  |  | Model statistics |  |  |  |
| --- | --- | --- | --- | --- | --- | --- | --- | --- |
| | | $\beta$ | t | p | R<sup>2</sup> | F | df | p |
| #1. SPT | Model |  |  |  | 0.140 | 6.455 | 2/79 | 0.003 |
|  | z FPG +z INS | 0.318 | 3.036 | 0.003 |  |  |  |  |
|  | Glucagon | -0.230 | -2.199 | 0.031 |  |  |  |  |
| #2. BW | Model |  |  |  | 0.799 | 103.571 | 3/78 | < 0.001 |
|  | Sex | 0.872 | 15.551 | < 0.001 |  |  |  |  |
|  | z FPG +z INS | 0.207 | 4.004 | < 0.001 |  |  |  |  |
|  | PAI-1 | 0.119 | 2.151 | 0.035 |  |  |  |  |
| #3. FI | Model |  |  |  | 0.375 | 23.675 | 2/79 | < 0.001 |
|  | Sex | 0.505 | 5.182 | < 0.001 |  |  |  |  |
|  | Leptin | 0.195 | 2.004 | 0.049 |  |  |  |  |
| #4. OFT surround | Model |  |  |  | 0.099 | 4.354 | 2/79 | 0.016 |
|  | GIP | -0.248 | -2.315 | 0.023 |  |  |  |  |
|  | Leptin | -0.215 | -2.003 | 0.049 |  |  |  |  |
| #5. OFT rearing | Model |  |  |  | 0.104 | 9.248 | 1/80 | 0.003 |
|  | Glucose | 0.322 | 3.041 | 0.003 |  |  |  |  |
| #6. NOR Preference Index | Model |  |  |  | 0.111 | 4.824 | 2/77 | 0.011 |
|  | Glucagon | 0.233 | 2.159 | 0.034 |  |  |  |  |
|  | PAI-1 | 0.216 | 1.996 | 0.049 |  |  |  |  |
| #7. FST Immobility Time | Model |  |  |  | 0.057 | 4.797 | 1/80 | 0.031 |
|  | z FPG +z INS | -0.238 | -2.190 | 0.031 |  |  |  |  |

SPT: Sucrose preference test. BW: Body weight. FI: Food intake. OFT surround: distance travelled in the surrounding zone of the open field. OFT rearing : number of rearing episodes in the Open Field Test. NOR Preference Index : Novel Object Recognition Preference Index for the novel object [novel/(novel+familiar)×100%]. FST Immobility Time: Forced Swim Test Immobility time.

## 4. Discussion

### 4.1 Insulin Resistance and Sensitivity in CUMS

The first key finding is that, under metabolically uncompromised baseline conditions—defined by standard chow feeding, stable body weight, and absence of obesogenic dietary challenge—mice exposed to CUMS developed enhanced insulin sensitivity, as evidenced by concurrent reductions in fasting insulin, fasting glucose, and the insulin resistance index. This observation aligns with our prior clinical report of reduced insulin resistance in Chinese patients with major depressive disorder (MDD) who lacked comorbid metabolic syndrome (Luo et al., 2025).

Our findings contrast with reports of CUMS-induced insulin resistance or glucose intolerance (Liang et al., 2021; Rong et al., 2024), likely reflecting protocol and phenotyping differences. CUMS stress load varies substantially across laboratories (Antoniuk et al., 2019): moderate/shorter paradigms may induce compensatory adaptation, whereas prolonged or high-intensity stress can drive glucocorticoid excess and HPA-axis dysregulation, leading to metabolic decompensation and insulin resistance (Aboushaar & Serrano, 2024; Berbudi et al., 2025). Moreover, fasting indices capture basal tone and are not interchangeable with dynamic measures (clamp, OGTT) of insulin action under challenge (Park et al., 2021).

### 4.2 Metabolic Hormone Profile in CUMS

A second major finding is that fasting insulin and glucose were reduced without detectable changes in the metabolic hormones measured here, indicating a selective shift in basal glycemic–insulinemic tone with preserved broader endocrine homeostasis. The appetite-regulatory axis remained intact: leptin tracked stable body weight, and ghrelin did not change, suggesting no perceived energy deficit sufficient to reset hypothalamic energy-balance control (Kern et al., 2012; Kim et al., 2010). Similarly, unchanged fasting glucagon, GLP-1, and GIP suggest preserved basal islet–enteroendocrine communication, consistent with selective suppression of basal β-cell secretory tone rather than generalized disruption of islet–gut crosstalk.

The accompanying mild hypoglycemia likely reflects reduced hepatic glucose output and/or enhanced peripheral glucose utilization under chronic stress, rather than insulin excess. One plausible mechanism is autonomic modulation that fine-tunes basal β-cell activity and glucose regulation during sustained stress (Ahrén, 2000; Kiba, 2004). In support of a non-obesity–linked metabolic adaptation, circulating resistin and PAI-1—adipokines typically elevated with obesity-related adipose inflammation—remained unchanged in CUMS mice (Hosogai et al., 2007).

### 4.3 Mechanism-Specific Pharmacological Efficacy in CUMS

While fluoxetine ameliorated depressive-like behaviors, it had minimal effects on the metabolic hormones assessed. In contrast, other compounds produced distinct peripheral metabolic/inflammation-linked signatures. Simvastatin increased GLP-1, consistent with pleiotropic anti-inflammatory and endothelial-protective effects of mevalonate-pathway inhibition (Lahera et al., 2007); given GLP-1’s neuroprotective and neurotrophic actions, this gut-derived signal may contribute to simvastatin’s antidepressant-like effects (During et al., 2003). Curcumin elevated PAI-1 and modestly increased ghrelin, suggesting modulation of the stress–coagulation–tissue remodeling axis through multi-target actions (e.g., NF-κB and TGF-β/Smad pathways) (Yu et al., 2021; Zhang et al., 2019). As a context-dependent mediator, PAI-1 may reflect either pro-fibrotic signaling or active repair, underscoring the complexity of curcumin’s systemic effects (Ghosh & Vaughan, 2012).

Both SAMe and PDTC increased circulating ghrelin but diverged in their effects on resistin (SAMe increased resistin, whereas PDTC markedly suppressed it). As a universal methyl donor, SAMe may influence adipokine expression through broad epigenetic reprogramming and nutrient-sensing functions that couple metabolism to chromatin and autophagic regulation. In contrast, PDTC’s suppression of the adipocyte-derived, pro-inflammatory adipokine resistin provides evidence for a targeted anti-inflammatory action (Silswal et al., 2005). Importantly, none of the interventions reversed the persistent hypoinsulinemic state in CUMS.

### 4.4 Metabolic Predictors of depressive -like behaviors

The insulin resistance index explained 21.1% of the variance in sucrose preference and 12.6% in FST immobility, suggesting that relatively higher basal insulin levels within this hypoinsulinemic context are associated with better affective outcomes (Grillo et al., 2015; Kim et al., 2025). This is consistent with evidence that central insulin signaling supports synaptic plasticity, dampens microglial NF-κB–linked neuroinflammation, and modulates dopaminergic transmission in reward circuitry—processes relevant to SPT and FST performance (Gruber et al., 2023). Thus, relative preservation of insulin may help maintain hippocampal and nucleus accumbens insulin signaling, buffering against stress-induced anhedonia and despair (Grillo et al., 2015).

Leptin predicted food intake and was inversely associated with open-field exploration, consistent with region-specific leptin actions on anxiety and motivation (Figge-Schlensok et al., 2025; Kwon et al., 2016). Glucagon showed dissociated behavioral associations (lower sucrose preference but higher NOR), whereas GLP-1 showed no behavioral links; these associations may reflect indirect peripheral-to-central signaling via hepatic vagal afferents and/or systemic glucose dynamics, potentially dampening mesolimbic reward while supporting hippocampal memory consolidation (Abraham & Lam, 2016; LaPierre et al., 2014). PAI-1 (increased with curcumin) predicted body weight and cognition, consistent with roles in extracellular proteolysis and brain-derived neutrophic factor (BDNF) maturation under stress (Party et al., 2019).

### 4.5 Sex Differences in CUMS mice

The last core finding is a clear sexual dimorphism: females showed higher resistin, PAI-1, glucagon, and ghrelin than males, whereas males exhibited higher leptin and insulin. This pattern suggests a female bias toward energy mobilization (ghrelin and glucagon) and heightened inflammation-linked tone (resistin and PAI-1), whereas males display a more anabolic configuration favoring energy storage (leptin and insulin) (Tramunt et al., 2020).

These sex-specific endocrine signatures align with clinical reports of higher depression prevalence in women, who often show more pronounced appetite disturbances and immune-related comorbidities (Altemus et al., 2014; Schuch et al., 2014). Such dimorphisms likely reflect coordinated crosstalk among sex hormones (estrogens/androgens), the HPA axis, metabolic tissues, and the immune system (Klein & Flanagan, 2016; Oyola & Handa, 2017). Notably, leptin regulation differs across species: women typically exhibit higher circulating leptin than men independent of adiposity, whereas male rodents often show higher leptin under standard laboratory housing (e.g., 22°C), a sub-thermoneutral condition for rodents. This divergence likely reflects differences in neuroendocrine–metabolic organization and temperature-sensitive sympathetic regulation of adipose leptin expression (Mulet et al., 2003; Trayhurn et al., 1995). Moreover, even at thermoneutrality (e.g., 28°C), sex differences in circulating leptin can persist despite diminished mRNA differences, implicating additional sex-dependent mechanisms (e.g., post-transcriptional regulation, clearance, or binding interactions) in leptin homeostasis (Mulet et al., 2003).

Estrogen may enhance female sensitivity to feeding-related and inflammatory signals, whereas androgens, despite directly suppressing leptin production in adipocytes, are associated with greater visceral adiposity in male rodents (Klein & Flanagan, 2016; Mahboobifard et al., 2022). This fat depot is relatively resistant to lipolytic stimuli, which may indirectly help sustain higher circulating leptin under standard housing conditions (Jia et al., 2025; Moinat et al., 1995). Collectively, this sex-dependent physiological context may shape distinct peripheral phenotypes in response to the same chronic stressor.

## Limitations

This study has several limitations. First, the CUMS model captures mainly stress-induced aspects of depression and was performed in metabolically healthy rodents, limiting generalizability to patients with comorbid metabolic dysfunction. Second, reliance on fasting serum measures prevents evaluation of dynamic metabolic responses or tissue-specific insulin sensitivity. Third, although both sexes were included, the modest sample size (n = 12/group) limited power to detect higher-order interactions. Fourth, drug dosing and timing were based on the literature without pharmacokinetic validation in our paradigm, potentially missing optimal windows or doses. Fifth, the behavioral test order could introduce carryover effects, and manual scoring may allow observer bias. Finally, we did not include drug-only (non-stressed) control groups for fluoxetine, simvastatin, curcumin, SAMe, or PDTC, reflecting a trade-off between causal attribution and adherence to 3R principles (Replacement, reduction, and refinement).

## Conclusions

CUMS induced a selective hypoinsulinemic fasting profile, with lower insulin and glucose levels, consistent with enhanced basal insulin sensitivity and distinct from the pathological insulin resistance typical of metabolic syndrome. This pattern suggests a compartmentalized, energy-conserving adaptation that co-occurs with behavioral resilience without overt disruption of central appetite regulation or adipose inflammation-linked signaling. A composite insulin-resistance index predicted affective outcomes, supporting basal insulin sensitivity as a key metabolic correlate of mood resilience. Pharmacological interventions targeting monoaminergic, epigenetic, lipid-metabolic, or NF-κB–related pathways differentially altered depressive-like behaviors and selected circulating factors (e.g., GLP-1, PAI-1), yet none reversed the core hypoinsulinemic state. Finally, pronounced sexual dimorphism emerged, with females showing a more catabolic, inflammation-vigilant endocrine profile and males a more anabolic, storage-oriented configuration, suggesting sex-dependent metabolic vulnerability to chronic stress.

## Supporting information

Electronic Supplementary File

## Acknowledgements

Not Applicable

## Data Availability Statement

The dataset generated during this study will be available upon reasonable request from the corresponding author (MM) upon publication.

## References

Aboushaar, N., & Serrano, N. (2024). The mutually reinforcing dynamics between pain and stress: Mechanisms, impacts and management strategies. Frontiers in Pain Research, 5, 1445280. 10.3389/fpain.2024.1445280

Abraham, M. A., & Lam, T. K. T. (2016). Glucagon action in the brain. Diabetologia, 59(7), 1367–1371. 10.1007/s00125-016-3950-3

Ahrén, B. (2000). Autonomic regulation of islet hormone secretion—Implications for health and disease. Diabetologia, 43(4), 393–410. 10.1007/s001250051322

Altemus, M., Sarvaiya, N., & Neill Epperson, C. (2014). Sex differences in anxiety and depression clinical perspectives. Frontiers in Neuroendocrinology, 35(3), 320–330. 10.1016/j.yfrne.2014.05.004

Antoniuk, S., Bijata, M., Ponimaskin, E., & Wlodarczyk, J. (2019). Chronic unpredictable mild stress for modeling depression in rodents: Meta-analysis of model reliability. Neuroscience & Biobehavioral Reviews, 99, 101–116. 10.1016/j.neubiorev.2018.12.002

Berbudi, A., Khairani, S., & Tjahjadi, A. I. (2025). Interplay between insulin resistance and immune dysregulation in type 2 diabetes mellitus: Implications for therapeutic interventions. ImmunoTargets and Therapy, 14, 359–382. 10.2147/ITT.S499605

Chen, X., Zhao, P., Wang, W., Guo, L., & Pan, Q. (2024). The antidepressant effects of GLP-1 receptor agonists: A systematic review and meta-analysis. American Journal of Geriatric Psychiatry, 32(1), 117–127. 10.1016/j.jagp.2023.08.010

Chouchani, E. T., & Kajimura, S. (2019). Metabolic adaptation and maladaptation in adipose tissue. Nature Metabolism, 1(2), 189–200. 10.1038/s42255-018-0021-8

Dregan, A., Rayner, L., Davis, K. A. S., Bakolis, I., Arias De La Torre, J., Das-Munshi, J., Hatch, S. L., Stewart, R., & Hotopf, M. (2020). Associations between depression, arterial stiffness, and metabolic syndrome among adults in the UK biobank population study: A mediation analysis. JAMA Psychiatry, 77(6), 598. 10.1001/jamapsychiatry.2019.4712

During, M. J., Cao, L., Zuzga, D. S., Francis, J. S., Fitzsimons, H. L., Jiao, X., Bland, R. J., Klugmann, M., Banks, W. A., Drucker, D. J., & Haile, C. N. (2003). Glucagon-like peptide-1 receptor is involved in learning and neuroprotection. Nature Medicine, 9(9), 1173–1179. 10.1038/nm919

Fan, C., Song, Q., Wang, P., Li, Y., Yang, M., Liu, B., & Yu, S. Y. (2018). Curcumin protects against chronic stress-induced dysregulation of neuroplasticity and depression-like behaviors via suppressing IL-1β pathway in rats. Neuroscience, 392, 92–106. 10.1016/j.neuroscience.2018.09.028

Fasshauer, M., & Blüher, M. (2015). Adipokines in health and disease. Trends in Pharmacological Sciences, 36(7), 461–470. 10.1016/j.tips.2015.04.014

Figge-Schlensok, R., Petzold, A., Hugger, N., Bakhareva, A., Abdallah, A. T., Wissing, C., Van Den Munkhof, H. E., Witt, M. Y., Awad, D. I., & Korotkova, T. (2025). A lateral hypothalamic neuronal population expressing leptin receptors counteracts anxiety to enable adaptive behavioral responses. Nature Neuroscience, 28(11), 2262–2272. 10.1038/s41593-025-02078-y

Gajewska, A., Strzelecki, D., & Gawlik-Kotelnicka, O. (2023). Ghrelin as a biomarker of “immunometabolic depression” and its connection with dysbiosis. Nutrients, 15(18), 3960. 10.3390/nu15183960

Ghosh, A. K., & Vaughan, D. E. (2012). PAI[1 in tissue fibrosis. Journal of Cellular Physiology, 227(2), 493–507. 10.1002/jcp.22783

Grillo, C. A., Piroli, G. G., Lawrence, R. C., Wrighten, S. A., Green, A. J., Wilson, S. P., Sakai, R. R., Kelly, S. J., Wilson, M. A., Mott, D. D., & Reagan, L. P. (2015). Hippocampal insulin resistance impairs spatial learning and synaptic plasticity. Diabetes, 64(11), 3927–3936. 10.2337/db15-0596

Gruber, J., Hanssen, R., Qubad, M., Bouzouina, A., Schack, V., Sochor, H., Schiweck, C., Aichholzer, M., Matura, S., Slattery, D. A., Zopf, Y., Borgland, S. L., Reif, A., & Thanarajah, S. E. (2023). Impact of insulin and insulin resistance on brain dopamine signalling and reward processing – An underexplored mechanism in the pathophysiology of depression? Neuroscience & Biobehavioral Reviews, 149, 105179. 10.1016/j.neubiorev.2023.105179

Haagsma, J. A., Graetz, N., Bolliger, I., Naghavi, M., Higashi, H., Mullany, E. C., Abera, S. F., Abraham, J. P., Adofo, K., Alsharif, U., Ameh, E. A., Ammar, W., Antonio, C. A. T., Barrero, L. H., Bekele, T., Bose, D., Brazinova, A., Catalá-López, F., Dandona, L., … Phillips, M. R. (2016). The global burden of injury: Incidence, mortality, disability-adjusted life years and time trends from the global burden of disease study 2013. Injury Prevention, 22(1), 3–18. 10.1136/injuryprev-2015-041616

Hasin, D. S., Sarvet, A. L., Meyers, J. L., Saha, T. D., Ruan, W. J., Stohl, M., & Grant, B. F. (2018). Epidemiology of adult *DSM-5* major depressive disorder and its specifiers in the United States. JAMA Psychiatry, 75(4), 336. 10.1001/jamapsychiatry.2017.4602

Hontecilla-Prieto, L., García-Domínguez, D. J., Berlanga-Gil, C., Flores-Campos, R., Muñoz-Pacheco, R., Franco-Fernández, M. D., Flores-Cordero, J. A., Sánchez-Jiménez, F., Pérez-Pérez, A., Vilariño-García, T., & Sánchez-Margalet, V. (2025). Leptin a potential link between obesity and depression. Cellular and Molecular Life Sciences, 82(1), 365. 10.1007/s00018-025-05892-6

Hosogai, N., Fukuhara, A., Oshima, K., Miyata, Y., Tanaka, S., Segawa, K., Furukawa, S., Tochino, Y., Komuro, R., Matsuda, M., & Shimomura, I. (2007). Adipose tissue hypoxia in obesity and its impact on adipocytokine dysregulation. Diabetes, 56(4), 901–911. 10.2337/db06-0911

Hotamisligil, G. S. (2017). Inflammation, metaflammation and immunometabolic disorders. Nature, 542(7640), 177–185. 10.1038/nature21363

Jia, Y., Lacombe, M., Muller, C., & Milhas, D. (2025). Adipose tissue and androgens: The ins and outs. Adipocyte, 14(1), 2508885. 10.1080/21623945.2025.2508885

Kern, A., Albarran-Zeckler, R., Walsh, H. E., & Smith, R. G. (2012). Apo-ghrelin receptor forms heteromers with DRD2 in hypothalamic neurons and is essential for anorexigenic effects of DRD2 agonism. Neuron, 73(2), 317–332. 10.1016/j.neuron.2011.10.038

Kiba, T. (2004). Relationships between the autonomic nervous system and the pancreas including regulation of regeneration and apoptosis: Recent developments. Pancreas, 29(2), e51–e58. 10.1097/00006676-200408000-00019

Kieffer, T. J., & Francis Habener, J. (1999). The glucagon-like peptides. Endocrine Reviews, 20(6), 876–913. 10.1210/edrv.20.6.0385

Kim, B., Kim, M., Lee, H.-Y., Pyo, J. H., Seo, J., Jeon, Y., Lee, H., Kim, J.-H., Ahn, S. H., Chi, S. W., Seong, J. K., & Baik, J.-H. (2025). Dopamine D2 receptor modulation of insulin receptor signaling in the central amygdala: Implications for compulsive-like eating behavior. Molecular Psychiatry, 0–12. 10.1038/s41380-025-03150-6

Kim, K. S., Yoon, Y. R., Lee, H. J., Yoon, S., Kim, S.-Y., Shin, S. W., An, J. J., Kim, M.-S., Choi, S.-Y., Sun, W., & Baik, J.-H. (2010). Enhanced hypothalamic leptin signaling in mice lacking dopamine D2 receptors. Journal of Biological Chemistry, 285(12), 8905–8917. 10.1074/jbc.M109.079590

Klein, S. L., & Flanagan, K. L. (2016). Sex differences in immune responses. Nature Reviews Immunology, 16(10), 626–638. 10.1038/nri.2016.90

Kubera, M., Obuchowicz, E., Goehler, L., Brzeszcz, J., & Maes, M. (2011). In animal models, psychosocial stress-induced (neuro)inflammation, apoptosis and reduced neurogenesis are associated to the onset of depression. Progress in Neuro-Psychopharmacology and Biological Psychiatry, 35(3), 744–759. 10.1016/j.pnpbp.2010.08.026

Kuehner, C. (2017). Why is depression more common among women than among men? The Lancet Psychiatry, 4(2), 146–158. 10.1016/S2215-0366(16)30263-2

Kwon, O., Kim, K. W., & Kim, M.-S. (2016). Leptin signalling pathways in hypothalamic neurons. Cellular and Molecular Life Sciences, 73(7), 1457–1477. 10.1007/s00018-016-2133-1

Lahera, V., Goicoechea, M., Garcia De Vinuesa, S., Miana, M., Heras, N. D. L., Cachofeiro, V., & Luno, J. (2007). Endothelial dysfunction, oxidative stress and inflammation in atherosclerosis: Beneficial effects of statins. Current Medicinal Chemistry, 14(2), 243–248. 10.2174/092986707779313381

LaPierre, M. P., Abraham, M. A., Filippi, B. M., Yue, J. T. Y., & Lam, T. K. T. (2014). Glucagon and lipid signaling in the hypothalamus. Mammalian Genome, 25(9–10), 434–441. 10.1007/s00335-014-9510-6

Liang, L., Zheng, Y., Xie, Y., Xiao, L., & Wang, G. (2021). Oridonin ameliorates insulin resistance partially through inhibition of inflammatory response in rats subjected to chronic unpredictable mild stress. International Immunopharmacology, 91, 107298. 10.1016/j.intimp.2020.107298

López-López, A. L., Jaime, H. B., Escobar Villanueva, M. D. C., Padilla, M. B., Palacios, G. V., & Aguilar, F. J. A. (2016). Chronic unpredictable mild stress generates oxidative stress and systemic inflammation in rats. Physiology & Behavior, 161, 15–23. 10.1016/j.physbeh.2016.03.017

Lu, S. C., & Mato, J. M. (2012). *S* -adenosylmethionine in liver health, injury, and cancer. Physiological Reviews, 92(4), 1515–1542. 10.1152/physrev.00047.2011

Luo, Y., Niu, M., Chen, T., Li, J., Almulla, A. F., Zhang, Y., & Maes, M. (2025). Lower insulin resistance in Chinese patients with severe major depressive disorder: Associations with the inflammatory response. medRxiv, 2025.10.10.25337709. 10.1101/2025.10.10.25337709

Maes, M., Almulla, A. F., You, Z., & Zhang, Y. (2025). Neuroimmune, metabolic and oxidative stress pathways in major depressive disorder. Nature Reviews Neurology, 21(9), 473–489. 10.1038/s41582-025-01116-4

Maes, M., Fišar, Z., Medina, M., Scapagnini, G., Nowak, G., & Berk, M. (2012). New drug targets in depression: Inflammatory, cell-mediated immune, oxidative and nitrosative stress, mitochondrial, antioxidant, and neuroprogressive pathways. And new drug candidates—Nrf2 activators and GSK-3 inhibitors. Inflammopharmacology, 20(3), 127–150. 10.1007/s10787-011-0111-7

Mahboobifard, F., Pourgholami, M. H., Jorjani, M., Dargahi, L., Amiri, M., Sadeghi, S., & Tehrani, F. R. (2022). Estrogen as a key regulator of energy homeostasis and metabolic health. Biomedicine & Pharmacotherapy, 156, 113808. 10.1016/j.biopha.2022.113808

Menze, E. T., Ezzat, H., Shawky, S., Sami, M., Selim, E. H., Ahmed, S., Maged, N., Nadeem, N., Eldash, S., & Michel, H. E. (2021). Simvastatin mitigates depressive-like behavior in ovariectomized rats: Possible role of NLRP3 inflammasome and estrogen receptors’ modulation. International Immunopharmacology, 95, 107582. 10.1016/j.intimp.2021.107582

Moinat, M., Deng, C., Muzzin, P., Assimacopoulos-Jeannet, F., Seydoux, J., Dulloo, A. G., & Giacobino, J.-P. (1995). Modulation of *obese* gene expression in rat brown and white adipose tissues. FEBS Letters, 373(2), 131–134. 10.1016/0014-5793(95)01030-I

Moreira, F. P., Jansen, K., Cardoso, T. D. A., Mondin, T. C., Magalhães, P. V. D. S., Kapczinski, F., Souza, L. D. D. M., Da Silva, R. A., Oses, J. P., & Wiener, C. D. (2017). Metabolic syndrome in subjects with bipolar disorder and major depressive disorder in a current depressive episode: Population-based study. Journal of Psychiatric Research, 92, 119–123. 10.1016/j.jpsychires.2017.03.025

Moreira, F. P., Jansen, K., Cardoso, T. D. A., Mondin, T. C., Vieira, I. S., Magalhães, P. V. D. S., Kapczinski, F., Souza, L. D. D. M., Da Silva, R. A., Oses, J. P., & Wiener, C. D. (2019). Metabolic syndrome, depression and anhedonia among young adults. Psychiatry Research, 271, 306–310. 10.1016/j.psychres.2018.08.009

Mulet, T., Picó, C., Oliver, P., & Palou, A. (2003). Blood leptin homeostasis: Sex-associated differences in circulating leptin levels in rats are independent of tissue leptin expression. International Journal of Biochemistry & Cell Biology, 35(1), 104–110. 10.1016/S1357-2725(02)00092-4

Olefsky, J. M., & Glass, C. K. (2010). Macrophages, inflammation, and insulin resistance. Annual Review of Physiology, 72(1), 219–246. 10.1146/annurev-physiol-021909-135846

Oyola, M. G., & Handa, R. J. (2017). Hypothalamic–pituitary–adrenal and hypothalamic–pituitary–gonadal axes: Sex differences in regulation of stress responsivity. Stress, 20(5), 476–494. 10.1080/10253890.2017.1369523

Park, S. Y., Gautier, J.-F., & Chon, S. (2021). Assessment of insulin secretion and insulin resistance in human. Diabetes & Metabolism Journal, 45(5), 641–654. 10.4093/dmj.2021.0220

Party, H., Dujarrier, C., Hébert, M., Lenoir, S., Martinez De Lizarrondo, S., Delépée, R., Fauchon, C., Bouton, M.-C., Obiang, P., Godefroy, O., Save, E., Lecardeur, L., Chabry, J., Vivien, D., & Agin, V. (2019). Plasminogen activator inhibitor-1 (PAI-1) deficiency predisposes to depression and resistance to treatments. Acta Neuropathologica Communications, 7(1), 153. 10.1186/s40478-019-0807-2

Rong, J., Wang, Y., Liu, N., Shen, L., Ma, Q., Wang, M., & Han, B. (2024). Chronic stress induces insulin resistance and enhances cognitive impairment in AD. Brain Research Bulletin, 217, 111083. 10.1016/j.brainresbull.2024.111083

Schuch, J. J. J., Roest, A. M., Nolen, W. A., Penninx, B. W. J. H., & De Jonge, P. (2014). Gender differences in major depressive disorder: Results from the Netherlands study of depression and anxiety. Journal of Affective Disorders, 156, 156–163. 10.1016/j.jad.2013.12.011

Sharma, S., Chawla, S., Kumar, P., Ahmad, R., & Kumar Verma, P. (2024). The chronic unpredictable mild stress (CUMS) paradigm: Bridging the gap in depression research from bench to bedside. Brain Research, 1843, 149123. 10.1016/j.brainres.2024.149123

Silswal, N., Singh, A. K., Aruna, B., Mukhopadhyay, S., Ghosh, S., & Ehtesham, N. Z. (2005). Human resistin stimulates the pro-inflammatory cytokines TNF-α and IL-12 in macrophages by NF-κB-dependent pathway. Biochemical and Biophysical Research Communications, 334(4), 1092–1101. 10.1016/j.bbrc.2005.06.202

Slavich, G. M., & Sacher, J. (2019). Stress, sex hormones, inflammation, and major depressive disorder: Extending social signal transduction theory of depression to account for sex differences in mood disorders. Psychopharmacology, 236(10), 3063–3079. 10.1007/s00213-019-05326-9

Tramunt, B., Smati, S., Grandgeorge, N., Lenfant, F., Arnal, J.-F., Montagner, A., & Gourdy, P. (2020). Sex differences in metabolic regulation and diabetes susceptibility. Diabetologia, 63(3), 453–461. 10.1007/s00125-019-05040-3

Trayhurn, P., Duncan, J. S., & Rayner, D. V. (1995). Acute cold-induced suppression of *ob* (obese) gene expression in white adipose tissue of mice: Mediation by the sympathetic system. Biochemical Journal, 311(3), 729–733. 10.1042/bj3110729

Yin, J., Wu, M., Duan, J., Liu, G., Cui, Z., Zheng, J., Chen, S., Ren, W., Deng, J., Tan, X., Al-Dhabi, N. A., Duraipandiyan, V., Liao, P., Li, T., & Yulong, Y. (2015). Pyrrolidine dithiocarbamate inhibits NF-KappaB activation and upregulates the expression of Gpx1, Gpx4, occludin, and ZO-1 in DSS-induced colitis. Applied Biochemistry and Biotechnology, 177(8), 1716–1728. 10.1007/s12010-015-1848-z

Yu, W.-K., Hwang, W.-L., Wang, Y.-C., Tsai, C.-C., & Wei, Y.-H. (2021). Curcumin suppresses TGF-β1-induced myofibroblast differentiation and attenuates angiogenic activity of orbital fibroblasts. International Journal of Molecular Sciences, 22(13), 6829. 10.3390/ijms22136829

Yuan, J., Pi, C., Shen, H., Zhou, B., Wei, Y., Dechsupa, N., & Zhao, L. (2025). Potential therapeutic benefits of curcumin in depression or anxiety induced by chronic diseases: A systematic review of mechanistic and clinical evidence. Frontiers in Pharmacology, 16, 1638645. 10.3389/fphar.2025.1638645

Zhang, J., Zheng, Y., Luo, Y., Du, Y., Zhang, X., & Fu, J. (2019). Curcumin inhibits LPS-induced neuroinflammation by promoting microglial M2 polarization via TREM2/ TLR4/ NF-κB pathways in BV2 cells. Molecular Immunology, 116, 29–37. 10.1016/j.molimm.2019.09.020

