## Supplementary material for "Chronic Unpredictable Stress Drives Non-pathological, Adaptive Metabolic Hormone Reprogramming with Reduced Insulin Resistance Linked to Depressive-like Behaviors": Electronic Supplementary File

**Electronic Supplementary File (ESF)**

ESF Figure 1 The experimental design and behavioral tests in each group.

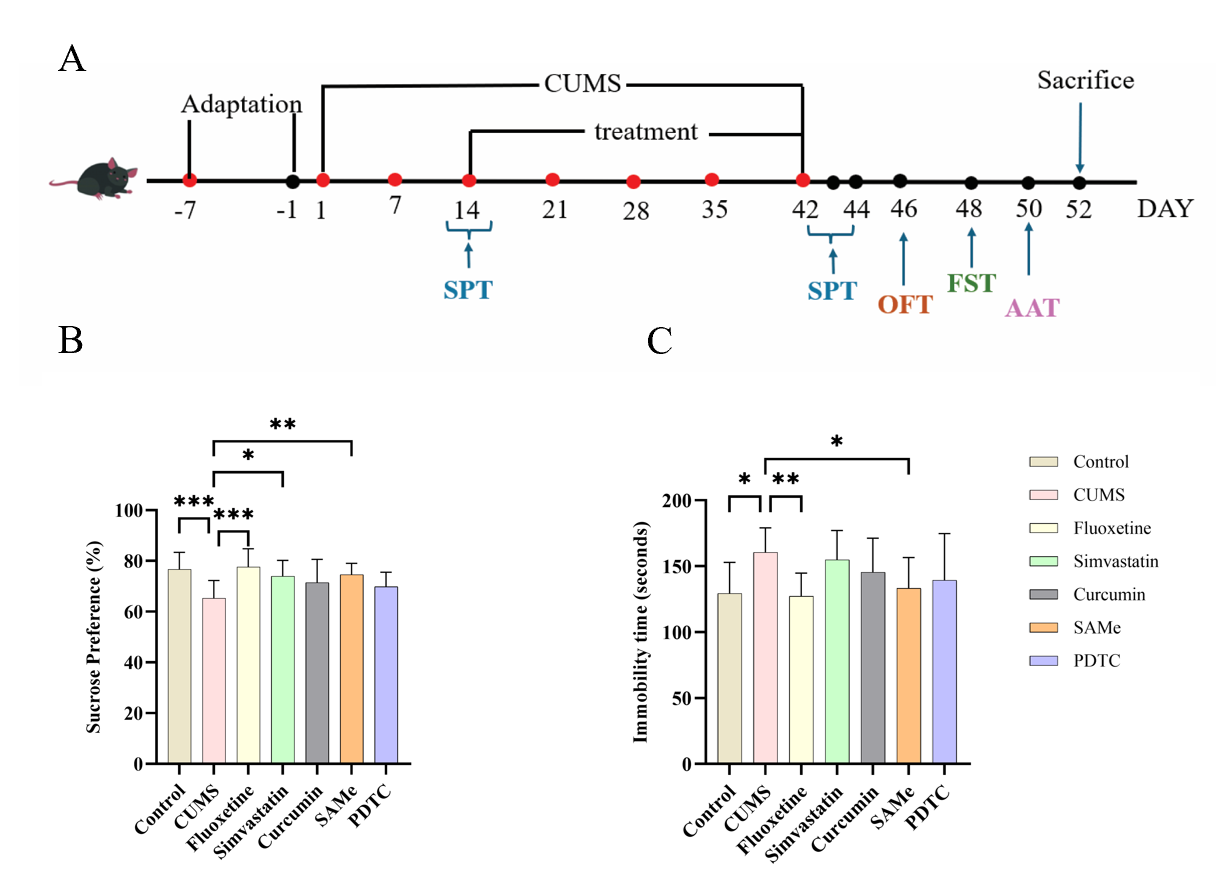

(A) Schematic timeline of the experimental procedure. Following a 7-day adaptation period in the facility (days -7 to -1), mice were randomly allocated to a control group (n=12, half male/female) or a chronic unpredictable mild stress (CUMS) modeling group (n=60, half male/female). After 2 weeks of CUMS exposure, a sucrose preference test (SPT) was conducted. CUMS mice were then subdivided into five treatment groups (n=12 each, half male/female): CUMS (vehicle), Fluoxetine, Simvastatin, Curcumin, and SAMe. CUMS induction and treatments continued for 4 additional weeks (days 14 to 42). Behavioral tests included SPT (day 42) and the forced swim test (FST, day 44). All mice were euthanized on day 47.
(B) Sucrose preference test (SPT) results across the seven groups: Control, CUMS, Fluoxetine, Simvastatin, Curcumin, SAMe, and PDTC.
(C) Forced swim test (FST) results across the seven groups: Control, CUMS, Fluoxetine, Simvastatin, Curcumin, SAMe, and PDTC.

ESF Table 1. Chronic unpredictable mild stress (CUMS) stressors and exposure parameters

| **Stressor domain** | **Stressor** | **Key parameters** | **Exposure duration** | **Schedule constraints** | **Source** |
| --- | --- | --- | --- | --- | --- |
| Sensory | Auditory/visual stimulation | 85 dB white noise + 3 Hz flashing light | 1 h | Weekly randomization; no consecutive-day repetition | Li et al., 2022 |
| Psychosocial | Social isolation | Single housing | 24 h | Weekly randomization; no consecutive-day repetition | Li et al., 2022 |
| Environmental | Cage tilt | 45° tilt | 24 h | Weekly randomization; no consecutive-day repetition | Li et al., 2022 |
| Deprivation | Water deprivation | Water removed | 24 h | Weekly randomization; no consecutive-day repetition | Li et al., 2022 |
| Environmental | Wet bedding | 200 mL water added to bedding | 24 h | Weekly randomization; no consecutive-day repetition | Li et al., 2022 |
| Physical | Restraint | Ventilated 50 mL tube | 2 h | Weekly randomization; no consecutive-day repetition | Li et al., 2022 |

Animals in the CUMS condition were exposed to two distinct mild stressors per day for 6 consecutive weeks. Stressors were randomized weekly, and the same stressor was not applied on consecutive days to minimize habituation. Control animals were housed under identical conditions without stress exposure.
Abbreviations: CUMS, chronic unpredictable mild stress.

ESF Table 2. Pharmacological interventions: formulations, dosing, and administration schedule

| **Group** | **Compound** | **Putative class / primary action** | **Supplier** | **CAS No.** | **Dose** | **Route** | **Vehicle / formulation** | **Treatment window** | **Citation** |
| --- | --- | --- | --- | --- | --- | --- | --- | --- | --- |
| Control + Vehicle | Vehicle control | — | — | — | — | Matched* | Matched* | q.d., weeks 3–6 (28 d) | — |
| CUMS + Vehicle | Vehicle control | — | — | — | — | Matched* | Matched* | q.d., weeks 3–6 (28 d) | — |
| CUMS + Fluoxetine | Fluoxetine | SSRI antidepressant | Aladdin (Shanghai, China) | 54910-89-3 | 20 mg/kg | Oral gavage | Saline | q.d., weeks 3–6 (28 d) | Li et al., 2022 |
| CUMS + Simvastatin | Simvastatin | Statin; pleiotropic immunometabolic effects | Aladdin (Shanghai, China) | 79902-63-9 | 10 mg/kg | Oral gavage | 1% DMSO | q.d., weeks 3–6 (28 d) | Yan et al., 2020 |
| CUMS + Curcumin | Curcumin | Polyphenol; antioxidant/anti-inflammatory | Aladdin (Shanghai, China) | 458-37-7 | 20 mg/kg | i.p. | Peanut oil (suspension) | q.d., weeks 3–6 (28 d) | — |
| CUMS + SAMe | S-adenosylmethionine (SAMe) | Methyl donor; metabolic/neuromodulatory | Solarbio (Beijing, China) | 97540-22-2 | 100 mg/kg | Oral gavage | Saline | q.d., weeks 3–6 (28 d) | Kulkarni et al., 2008 |
| CUMS + PDTC | Pyrrolidine dithiocarbamate (PDTC) | NF-κB pathway inhibitor | Aladdin (Shanghai, China) | 5108-96-3 | 140 mg/kg | Oral gavage | Saline | q.d., weeks 3–6 (28 d) | Qin et al., 2014 |

Starting 2 weeks after CUMS initiation (experimental week 3), mice were randomized into seven groups (n = 12/group, sex-balanced) and received treatments once daily for 28 days (weeks 3–6). All drug solutions/suspensions were freshly prepared under dim light. Vehicle-control procedures should match the corresponding administration route and vehicle used for each treatment condition.

Abbreviations: CUMS, chronic unpredictable mild stress; SAMe, S-adenosylmethionine; PDTC, pyrrolidine dithiocarbamate; SSRI, selective serotonin reuptake inhibitor; i.p., intraperitoneal; DMSO, dimethyl sulfoxide; q.d., once daily.

***ESF Methods***

*ESF Methods S1. Chronic unpredictable mild stress (CUMS) procedure*

The chronic unpredictable mild stress (CUMS) paradigm was conducted for 6 consecutive weeks. Animals assigned to the CUMS condition received two different mild stressors per day, randomly selected from the stressor set described in Table S1. Stressor sequences were re-randomized weekly, and the same stressor was not applied on two consecutive days to minimize habituation. Control animals were housed under identical environmental conditions without stress exposure.

*ESF Methods S2. Pharmacological interventions: preparation and administration*

Drug treatments began 2 weeks after CUMS initiation (experimental week 3) and were administered once daily for 28 days (weeks 3–6). All formulations were freshly prepared under dim light. Drug identity, supplier, CAS number, dose, route of administration, and vehicle/formulation are summarized in Table S2.

Vehicle control. Vehicle-treated control animals received the corresponding vehicle using the same administration route and dosing schedule as the drug-treated groups, as appropriate for the experimental design.

*ESF Methods S3. Body weight/food intake monitoring and behavioral procedures*

*ESF Methods S3.1.* Body weight and food intake

Body weight (BW) and food intake (FI) were recorded weekly throughout the experiment.

*ESF Methods S3.2.* General behavioral testing conditions

Behavioral tests were conducted during experimental week 7 in the following order to reduce carryover effects: Open Field Test (OFT) → Sucrose Preference Test (SPT) → Novel Object Recognition (NOR) → Forced Swim Test (FST). Experimenters responsible for testing and scoring were blinded to group assignment.

*ESF Methods S3.3. Open Field Test (OFT)*

To assess locomotor activity and anxiety-like behavior, mice were placed in the center of a square open-field arena (50 × 50 cm) under approximately 50 lux illumination and allowed to explore for 10 min. Locomotion (total distance traveled) and time/activity in the center zone (25 × 25 cm) were recorded and analyzed using an automated tracking system (RWD system).

*ESF Methods S3.4. Sucrose Preference Test (SPT)*

To assess anhedonia-like behavior, mice were habituated to two-bottle choice (1% sucrose solution vs. water) for 48 h. After 6 h food and water deprivation, intake from each bottle was measured over 12 h with bottle positions counterbalanced. Sucrose preference was calculated as:

$$\text{Sucrose preference (\%)}=\frac{\text{sucrose intake}}{\text{sucrose intake + water intake}}\times100$$

*ESF Methods S3.5. Novel Object Recognition (NOR)*

To assess recognition memory, mice were first allowed to explore two identical objects (training phase) for 10 min. Twenty-four hours later, one object was replaced with a novel object (different shape/color/texture; similar size), and mice explored for 10 min. The recognition index was calculated as:

$$\text{Recognition index (\%)}=\frac{\text{time exploring novel}}{\text{time exploring novel + familiar}}\times100$$

Objects and arenas were cleaned with 75% ethanol between trials. Exploratory behavior was defined as the mouse’s nose being within approximately 2 cm of the object.

*ESF Methods S3.6. Forced Swim Test (FST)*

To assess behavioral despair-like behavior, mice were placed individually in a transparent cylinder (10 cm diameter) filled with water (25 ± 1°C, 20 cm depth) for 6 min. Immobility time was scored during the last 4 min, with immobility defined as minimal movements necessary to keep the head above water.

*ESF Methods S4. Blood collection and biochemical assays*

*ESF Methods S4.1. Serum collection and storage*

At least 24 h after the final behavioral test (experimental day 47), mice were fasted overnight (approximately 16 h) and euthanized by cervical dislocation. Whole blood was collected via cardiac puncture. Serum was separated by centrifugation (3500 rpm, 10 min, 4°C), aliquoted immediately, and stored at −80°C until batch analysis.

*ESF Methods S4.2. Multiplex hormone quantification (Luminex/Bio-Plex)*

Serum levels of eight metabolic hormones (ghrelin, GIP, GLP-1, glucagon, insulin, leptin, PAI-1, resistin) were quantified using the Bio-Plex Pro™ Mouse Diabetes 8-Plex Assay (Bio-Rad; #171F7001M) based on Luminex® xMAP® magnetic bead technology. Serum samples were diluted 1:4 in the kit-provided Sample Diluent. Assays were performed according to the manufacturer’s protocol, including incubation with antibody-conjugated beads (1 h), incubation with biotinylated detection antibodies (30 min), and incubation with streptavidin-phycoerythrin (SA-PE; 10 min) on a microplate shaker (850 rpm) at room temperature. Fluorescence signals were acquired on a Luminex 200™ system, and concentrations were calculated from standard curves using Bio-Plex Manager™ software. All samples were run in technical duplicate.

*ESF Methods S4.3. Fasting serum glucose measurement*

Fasting serum glucose was measured colorimetrically using the o-toluidine method (Glucose Assay Kit, Beyotime Biotechnology; Cat# S0201S). Briefly, 5 μL serum was mixed with 185 μL reagent, heated at 95°C for 8 min, and absorbance was measured at 630 nm.

*ESF Methods S5. Statistical analysis details*

*ESF Methods S5.1. General principles*

All analyses were conducted in SPSS 27.0 using two-tailed tests with α = 0.05. One control female mouse died and was excluded; all remaining observations were complete.

*ESF Methods S5.2. Derived composite index*

A composite fasting glycemic/insulinemic burden index was computed as the sum of standardized scores:

$$\text{Composite index}=z(\text{FPG})+z(\text{INS})$$

where z-scores were computed as $(x-\mu)/\sigma$within the analyzed dataset.

*ESF Methods S5.3. Group comparisons*

For metabolic biomarker panels, sex was included as a fixed factor in all models. Group differences were evaluated using MANCOVA with fixed factors including treatment and sex. Behavioral outcomes were analyzed using unpaired t-tests for two-group comparisons or one-way ANOVA for multi-group comparisons, with appropriate post hoc tests as implemented in SPSS.

*ESF Methods S5.4. Exploratory biomarker–behavior associations*

To identify biomarkers independently associated with behavioral outcomes, stepwise multiple linear regression was applied exploratorily (entry criterion p < 0.05; removal criterion p > 0.10). Regression results were reported using standardized β coefficients, 95% confidence intervals, F-statistics, and R².

References:

Li, Q., Zhao, B., Li, W., He, Y., Tang, X., Zhang, T., Zhong, Z., Pan, Q., & Zhang, Y. (2022). Effects of repeated drug administration on behaviors in normal mice and fluoxetine efficacy in chronic unpredictable mild stress mice. *Biochemical and Biophysical Research Communications*, *615*, 36–42. https://doi.org/10.1016/j.bbrc.2022.05.041

Yan, J., Liu, A., Fan, H., Qiao, L., Wu, J., Shen, M., Lai, X., & Huang, J. (2020). Simvastatin improves behavioral disorders and hippocampal inflammatory reaction by NMDA-mediated anti-inflammatory function in MPTP-treated mice. *Cellular and Molecular Neurobiology*, *40*(7), 1155–1164. https://doi.org/10.1007/s10571-020-00804-7

Kulkarni, S. K., Bhutani, M. K., & Bishnoi, M. (2008). Antidepressant activity of curcumin: Involvement of serotonin and dopamine system. *Psychopharmacology*, *201*(3), 435–442. https://doi.org/10.1007/s00213-008-1300-y

Qin, J., Cao, Z., Li, X., Kang, X., Xue, Y., Li, Y., Zhang, D., Liu, X.-Y., & Xue, Y. (2014). Effect of ammonium pyrrolidine dithiocarbamate (PDTC) on NF-κB activation and CYP2E1 content of rats with immunological liver injury. *Pharmaceutical Biology*, *52*(11), 1460–1466. https://doi.org/10.3109/13880209.2014.898075
